# Capsinoids (non-pungent TRPV1 agonists) promote mild hypothermia and improved outcome following stroke in aged mice

**DOI:** 10.64898/2026.06.05.730536

**Authors:** Alexander P. Andersohn, Sodam Kim, Ting Wu, Andrea N. Doan, Charles L. Cantrell, Robert L. Jarret, Gab S. Kim, Sean P. Marrelli

**Affiliations:** Department of Neurology, University of Texas McGovern Medical School at Houston, Houston, TX, USA; Natural Products Utilization Research Unit, Agricultural Research Service, United States Department of Agriculture, University, MS, 38677, USA; Plant Genetic Resources Conservation Unit, Agricultural Research Service, United States Department of Agriculture, 1109 Experiment Street, Griffin, GA 30223, USA

**Keywords:** Capsinoids, Capsiate, Dihydrocapsiate, Transient Receptor Potential Vanilloid 1, Therapeutic Hypothermia, Ischemic Stroke, Neuroprotection

## Abstract

**Background and Purpose:** Mild hypothermia provides potent neuroprotection in experimental ischemic stroke, however, its implementation in awake stroke patients is hampered by the induction of intense shivering and inconsistent body temperature control. Pharmacological activation of peripheral/peritoneal TRPV1 channels with non-pungent capsinoids offers a means to induce hypothermia while minimizing TRPV1 activation in the injury region. We tested whether capsinoid-mediated mild hypothermia, initiated within the post-stroke period, reduces brain injury and improves functional outcomes in aged mice.

**Methods:** Aged (18–20 months) male and female C57BL/6 mice underwent either permanent distal middle cerebral artery (MCA) occlusion (pdMCAO) or 60-minute MCA occlusion/reperfusion (MCAO/R). At 2 or 4 hours after stroke, mice received intraperitoneal injections of vehicle or capsinoids (>97% purity; 40 mg/kg) every 90 minutes to induce mild hypothermia for 4.5–6 hours. Core temperature was monitored by wireless probe. After pdMCAO, infarct volume was quantified at post-stroke day 3 (PSD3) by TTC and brain atrophy at PSD30 by iodine-enhanced microCT; sensorimotor function (DigiGait, forelimb grip strength, foot fault) was assessed at PSD7 and PSD30. Survival was the primary measured outcome for MCAO/R.

**Results:** Capsinoids induced a rapid and sustained reduction in core temperature of 2–4°C, independent of sex. In the pdMCAO model, capsinoid-induced hypothermia reduced infarct volume by 48% at PSD3 and decreased chronic cortical tissue loss by 44% at PSD30. Capsinoid-treated mice showed significant improvements in gait, grip strength, and contralateral foot fault performance at PSD7 and PSD30. In the MCAO/R model, survival was significantly higher in capsinoid-treated mice (80%) versus vehicle controls (33%) through PSD3.

**Conclusions:** Intraperitoneal capsinoid administration after stroke induces mild hypothermia in aged mice and confers robust acute neuroprotection and improved chronic functional outcome and survival. These preclinical findings add support for the use of capsinoids as a means to target peripheral thermoeffectors for promoting neuroprotective hypothermia in conscious stroke subjects.

## Introduction

One in four people over 25 years of age are expected to experience a stroke in their lifetime[1]. Treatment options for acute stroke currently include thrombolysis and thrombectomy. While each of these treatment options can be very effective, multiple factors can limit their effectiveness or timely availability to patients. Guidelines for thrombolytic agents limits their use to within 4.5 hours of known stroke onset [2].

Thrombectomy offers a treatment window of up to 24 hours, however, its application is limited to a subset of patients based on the clot location being within a large vessel, imaging revealing evidence of salvageable brain tissue, and the availability of an endovascular interventional team[2]. As a result, a significant number of patients are effectively left without treatment for acute ischemic stroke [3–14]. Furthermore, even for patients that receive one of these interventions, fewer than half demonstrate a good outcome, such as a modified Rankin score of 2 or better[3–14]. Thus, there is still great need for improved therapeutic approaches.

Therapeutic hypothermia (TH) has shown clinical benefit with certain ischemic injuries, such as witnessed cardiac arrest and neonatal hypoxia [15–24]. For ischemic stroke, application of hypothermia has been shown to be very effective in experimental stroke models [25–33]. However, in conscious stroke patients, the approach is significantly complicated by the logistics of cooling awake patients [25, 34–38]. Current clinical approaches to therapeutic hypothermia involve external forced cooling or intravascular cooling via intravenous infusion of cold saline or insertion of a heat exchange device. These methods can be effective in lowering core body temperature, but often introduce clinical complications such as profound shivering, cardiac arrhythmia, and hypotension [26, 32]. To counter the effects of activating the body’s cold defense mechanism, subjects typically require strong sedatives or even paralytics [25, 34–38]. These caveats make existing cooling methods either insufficient or unsuitable for conscious stroke patients.

We and others have applied pharmacological approaches to induce a drop in whole-body temperature (Tb) by selectively targeting the TRPV1 channel within the thermoregulatory system [31, 32, 39–41]. The TRPV1 channel is activated by warm temperature (>43 °C) as well as by capsaicinoids (e.g. capsaicin, dihydrocapsaicin) [42–45]. By pharmacologically activating the TRPV1 channels within the thermoregulatory system, a false peripheral overheating signaling is relayed to the brain and a whole body cooling mechanism is activated [31–33, 40, 41, 46, 47]. Until recently, the TRPV1 agonists used to promote body cooling in these studies have been largely limited to the capsaicinoids. However, capsaicinoids can distribute throughout the body and are associated with activation of pain pathways. Our recent work showed that capsinoids, which are known as ‘non-pungent’ TRPV1 agonists, can also promote a drop in Tb in conscious mice [46]. Unlike capsaicinoids, the capsinoids are vulnerable to enzymatic breakdown by ubiquitous esterases, thus effectively limiting their site of action (i.e. TRPV1 channel activation) to the region of delivery [46]. We have exploited this molecular characteristic to promote TRPV1 activation selectively within the peritoneal cavity and activation of TRPV1-containing afferents within the thermoregulatory system. With repeated capsinoid delivery, mild hypothermia could be induced and maintained for more than 6 hours in adult and aged mice of both sexes [46]. Here, we tested the overall hypothesis that capsinoid-mediated mild hypothermia initiated 2 hours after stroke can provide neuroprotection and improved functional recovery in mice. To enhance translational value of this study, experiments were performed in aged mice (18-20 months) of both sexes.

## Materials and Methods

### Animals

Experiments were performed with aged (18-20 months) C57BL6 male and female mice. The mice were obtained from the NIA aging colony and housed for at least one month locally before experimental use. 29 total mice (15 males, 14 females) were used in the permanent acute stroke study, and 16 mice (8 males, 8 females) for the permanent chronic stroke study. While the permanent stroke model has a very high survival rate, one of the male mice in the vehicle group did not survive to the 30 day experimental endpoint. For the transient stroke study, 22 mice (11 males, 11 females) were used, with 10 mice failing to reach the 3 day experimental endpoint (5 males in the vehicle group, 5 females in the vehicle group, 1 male in the capsinoid group, and 1 female in the capsinoid group). Euthanasia for all experiments was carried out in the following manner: mice were deeply anesthetized with 2.5% avertin (1% by body weight, IP) and then perfused transcardially with heparinized PBS (10 U/mL) followed by 4% paraformaldehyde (PFA). All animal studies were approved by the Institutional Animal Care and Use Committee at the University of Texas Health Science Center at Houston.

### Temperature Probe Implant and Recording

Two days prior to any planned experiments, mice were briefly anesthetized with isoflurane (3-4% induction in 70%/30% oxygen/nitrogen) to implant temperature probes. Miniature wireless Implantable Programmable Temperature Transponder (IPTT-300, BioMedic Data Systems, Seaford, DE) were implanted beneath the skin behind the right scapula. Temperatures were read by a wireless reader and then logged. All recordings began with at least 30 minutes of baseline prior to first drug administration and recorded every 10 minutes. All temperature transponders were given a unique, non-group identifying moniker to allow for unbiased blinded data analysis.

### Capsinoid Extraction

Capsinoids were isolated from *Capsicum* species fruit by first using a non-polar organic solvent (pentane), second liquid/liquid partitioning using acetonitrile (ACN)/pentane, and lastly a step gradient C-4 purification using H_2_O/ACN as the mobile phase [48]. This yields a >97% purity ACN fraction consisting of capsiate (56%) and dihydrocapsiate (41%) measured by high performance liquid chromatography (HPLC).

### Intraperitoneal Injection (IP)

Vehicle (corn oil) or capsinoids, were injected into the intraperitoneal cavity of mice at a maximum volume of 1% of the mouse’s body weight. The injections were administered at a concentration of 40mg/kg. For repeated injections, the side of injection was swapped with each administration.

### Permanent Distal Middle Cerebral Artery Occlusion (pdMCAO) Model

The pdMCAO surgery was performed identically to what has been described previously. Briefly, mice were anesthetized with isoflurane (4% induction and 1.75-2% maintenance) and body temperature was maintained at 37°C via rectal probe and a feedback-controlled heating pad. Using a micro-drill, a small hole was generated in the skull just above the distal middle cerebral artery (MCA). A low temperature cautery was used to permanently ligate the MCA. During recovery, the mice were placed in a custom-built warming chamber [33] for two hours to maintain normothermic body temperature. Over the last 30 minutes of recovery, the core body temperature of the mice was measured by the implanted temperature probe and registered as baseline. The mice were then removed from the recovery chambers and administered 40mg/kg capsinoids or vehicle control at two hours after stroke induction (time 0) and every 90 minutes thereafter to provide 6 hours of hypothermia or normothermia. During this time, temperatures were recorded every 10 minutes. Mice were monitored daily for 3 days after surgery (for all mice) and then every other day until day 7 and spot checked as necessary until day 30 (in the 1 month cohort).

### Triphenyltetrazolium Chloride (TTC) Infarct Analysis

At the conclusion of the 3 day pdMCAO experiments a lethal injection of avertin was administered and the mice were perfused with PBS mixed with heparin. The brains were then removed and sectioned into 1mm slices, then immersed in 3% 2,3,5-triphenyltetrazolium chloride (TTC) solution in PBS at 37°C for 5 minutes, at which point they were flipped and stained for another 5 minutes. TTC is converted into a red compound by active mitochondria, thereby staining metabolically active tissue red and leaving infarcted tissue white. The slices were then fixed in 4% paraformaldehyde and quantified using ImageJ software (National Institutes of Health, Bethesda, MD). Each slice was imaged from both sides, length calibrated, total area calculated by “polygon selections” around the whole slice, infarct area calculated by “polygon selections” around infarct core and penumbra damage combined, and volumes calculated by areas multiplied by half-depth per image (0.5mm) and summed across all slices.

### Middle Cerebral Artery Occlusion with Reperfusion (MCAO/R)

Aged (18-20m) C57BL6 mice of both sexes were anesthetized with isoflurane (4% induction and 1.75-2% maintenance) and temperature was maintained at 37°C. The common, external, and internal carotid arteries were isolated and a small incision was made in the external carotid to insert and guide a silicone-coated monofilament into the right internal carotid artery and through to the Circle of Willis to block the middle cerebral artery. The mice were allowed to awaken after occlusion in a warming cage and neurological deficits were evaluated to confirm successful stroke. After 60 minutes, mice were re-anesthetized and the filament was removed to allow reperfusion. Mice were then returned to the warming cage for 3 hours before experimental treatment was initiated. Mice were monitored daily for 3 days after surgery and euthanized if it was determined that humane end points were displayed. At 3 days post-stroke, the mice were euthanized, perfused, and stained with TTC as described above.

### MicroCT Imaging and Analysis

1 month after pdMCAO, mice were given a lethal dose of avertin and transcardially perfused with PBS/heparin followed by PFA. Brains were removed and submerged in PFA overnight. The following day, the brains were submerged in 1% iodine by weight (Sigma Aldrich 229695) in 90% methanol for 3 days, replacing the solution after the first day. On the day of imaging, the brains were washed in distilled water for 1 hour and then embedded in 1% low melt agarose in PBS in a 5mL plastic vial. The brains were then imaged on a SkyScan 1276 CMOS Edition (MicroCT Imaging Core at the McGovern Medical School at UTHealth). The parameters for scanning were as follows: source voltage: 73kV, source current: 200uA, image pixel size: 5um, filter: 1mm Al, 4096 pixels x 4096 pixels, exposure: 365ms, averaged across 3 images, 3600 images over 360°. Acquired images were then reconstructed using NRecon Reconstruction software (Bruker Micro CT, Kontich, Belgium) and then rendered into quantifiable 3D datasets in Dragonfly volume rendering software (Comet, Switzerland). Ipsilateral and contralateral volumes were quantified across 600 coronal slices and the infarcted volume to total volume percentage was calculated.

### DigiGait Imaging and Analysis

DigiGait (MouseSpecifics, Inc) uses a transparent treadmill belt and a high speed camera positioned under the belt in order to capture a mouse’s gait pattern. The treadmill was set to a speed of 15 cm/s and videos were recorded until 10 uninterrupted successive steps were captured. The videos were then analyzed using the DigiGait Analysis 15 software.

*Grip Strength Test*.

Mice body weight was recorded before testing. The mice were then suspended by their tails above a wire mesh attached to potentiometer device that would record maximal grasping force before being lowered until just the fore paws could make contact. The mice would then be gently pulled away until they lose their grip and the maximal grasping strength would be recorded in grams. The grip strength would then be divided by the body weight in order to get a value that is more consistent between sexes and sizes of mice. The test was done in triplicate and the average was taken for each subject.

### Foot Fault Test

Mice were placed on a series of parallel rods that are separated by 1 cm. A GoPro camera is placed below the metal rods to record the underside of the mice. The mice were then left to freely wander on the apparatus in a quiet room for 2 minutes. The videos were then analyzed to count the number of times each paw fell through the rods as well as the total number of steps across the recorded 2 minutes. The total number of times the contralateral paws (both fore and hind) dropped below the plane of the metal rods (contralateral foot faults), the total number of times all paws dropped below the plane of the metal rods (total foot faults), and the total number of steps taken were all recorded for analysis.

### Data Presentation and Statistical Analysis

All data is presented as mean ± standard error of the mean (SEM). All temperature is represented as delta temperature (ΔT_b_ °C), defined as the difference between every datum and the baseline temperature measurement’s average. Representing the temperature in this manner limits potential complications and variability due to transponder-to-transponder differences as well as potential effects of minor differences in implantation sites. For comparative data, two-way repeated ANOVA was used for comparing the breadth of temperature responses, and where appropriate, individual time point differences. For column comparisons, Welch’s two-tailed t-test was performed for statistical significance.

## Results

All mice in this group underwent the permanent distal middle cerebral artery occlusion (pdMCAO) procedure and were then immediately transferred to a custom heat support cage to maintain post-stroke Tb within the normothermic range for two hours [33]. Mice regained consciousness and were ambulatory shortly after transfer to the heat support cage. Within the final 30 minutes of the heat support period, measured Tb was similar for both groups and within the normothermic range (Capsinoid: 36.7±0.1°C, Vehicle: 36.2±0.2°C, n=16/13). At 2 hours post-stroke, heat support was removed and mice were given either vehicle or capsinoids (40mg/kg; i.p.) to induce mild hypothermia (**Fig 1B**).

**Figure 1:**
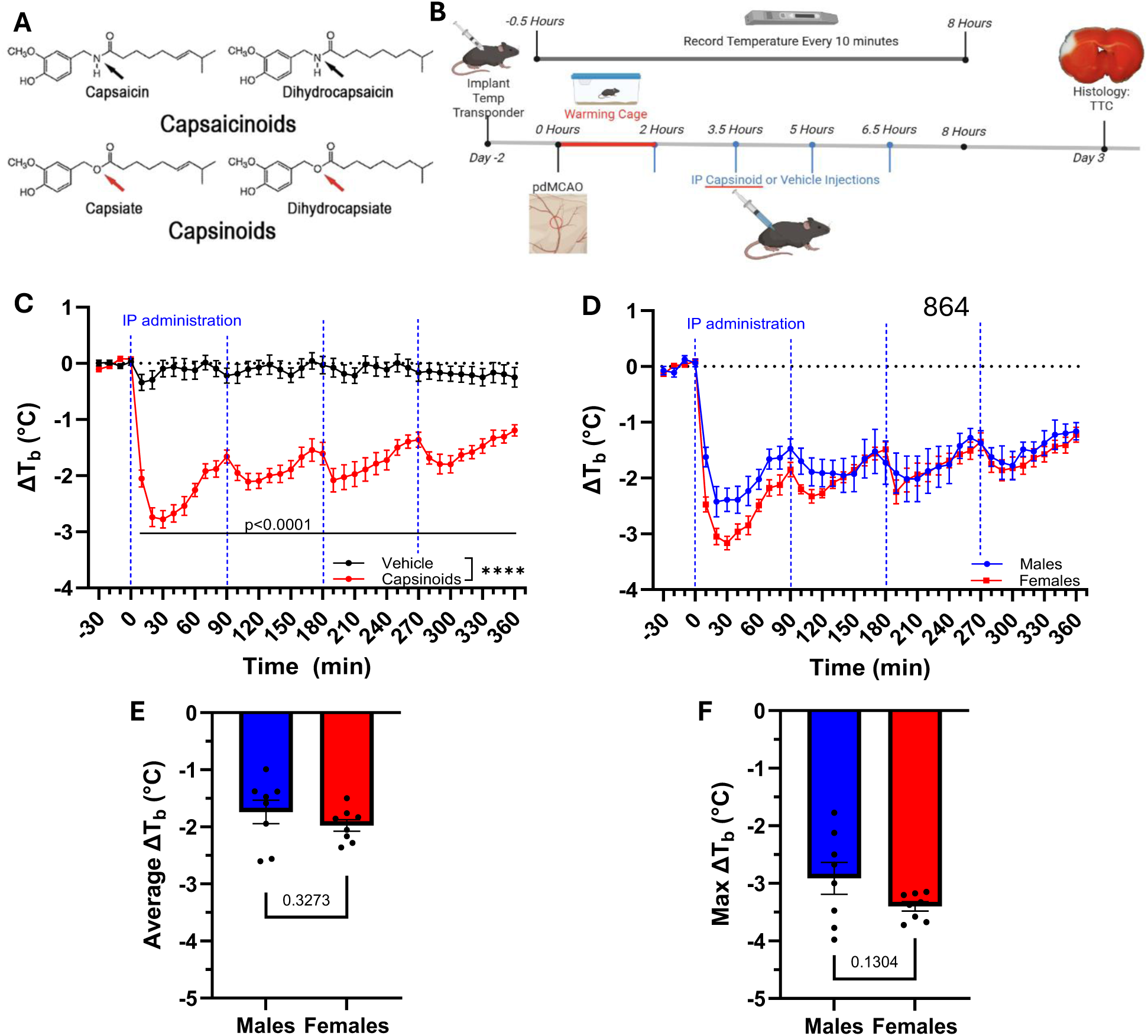
Intraperitoneally administered capsinoids induce a mild hypothermia after pdMCAO stroke. **A:** Chemical structures of capsaicinoids and capsinoids. Black arrows represent amide groups and red arrows represent ester groups. **B:** Experimental design showing timing of transponder implantation, pdMCAO stroke, recovery time, IP administration of vehicle or capsinoids, recording time, and experimental endpoints. **C:** Wireless measurement of ΔTb in aged mice beginning 2 hours after pdMCAO stroke with repeated IP administration of vehicle or capsinoids (40 mg/kg; indicated by blue dashed vertical lines). Data are presented as the combined group of males and females. **D:** Sex-specific comparison data from panel C. **E-F:** Summary of the female and male responses to repeated IP capsinoid administration compared as average ΔTb after administration (0-420 min) (**e**) and max drop in Tb (**f**). (*: p<0.05, **: p<0.01, ***: p<0.001, #: p<0.0001, or listed as numbers)

Delivery of capsinoids promoted a rapid initial drop in Tb, which was sustained for several hours with repeated injections at 90 min interval (**Fig 1C**). Vehicle treated mice demonstrated little change in Tb and were able to maintain normothermic Tb in the absence of heat support. The Tb change from baseline at 30 minutes post injection was -2.7±0.2 °C for capsinoids, versus 0.1±0.1 °C for vehicle (two-way RM measures ANOVA, p<0.0001, n=16/13). As stroke is a sexually dimorphic disease [49–51], we further compared the capsinoid-induced drop in Tb by sex (**Fig 1D**). Similar to what we previously found in naïve mice [46], the post-stroke mice demonstrated no statistical difference in the hypothermic effect when comparing the average Tb response (**Fig 1E**, M: -1.7±0.2°C, F: -2.0±0.1°C, two-tailed t-test, p=0.3273, n=8 per sex) or maximal drop in Tb (**Fig 1F**, M: -2.7±0.3°C, F: -3.4±0.1°C, two-tailed t-test, p=0.1304, n=8 per sex).

We first determined the effect of capsinoid-induced hypothermia on infarct volume in the acute stroke phase with the pdMCAO model. Infarct volume was evaluated by TTC staining of serial sections at post-stroke day (PSD) 3 (**Fig 2A**). Capsinoid-treated mice demonstrated a significant reduction in infarct volume compared to vehicle-treated mice (11.4±1.3 mm^3^ vs. 21.8±2.2 mm^3^, two-tailed t-test, p=0.0007, n=16/13) (**Fig 2B**). When normalizing infarct as a percent of total volume (IV/TV * 100), capsinoid treatment similarly showed significant reduction of infarct (2.6±0.3% vs. 5.3±0.5% (two-tailed t-test, p=0.0003, n=16/13) (**Fig 2C**). When separately analyzed by sex, capsinoid treatment reduced infarct volume in both males and females (**Fig 2D-G**). Capsinoid treatment reduced total infarct volume by 56.9% in males (8.7±1.4 mm^3^ (capsinoids) vs. 20.2±3.9 mm^3^ (vehicle); two-tailed t-test, p=0.0116, n=8/7) and by 41.1% in females (14.0±1.9 mm^3^ (capsinoids) vs. 23.8±1.9 mm^3^ (vehicle); two-tailed t-test, p=0.0033, n=8/6). Evaluated as percent infarct, capsinoids treatment reduced infarct by 59.6% in males (1.9±0.3% (capsinoids) vs. 4.7±0.8% (vehicle); two-tailed t-test, p=0.0054, n=8/7) and by 42.3% in females (3.4±0.5% (capsinoids) vs. 6.0±0.5% (vehicle); two-tailed t-test, p=0.0024, n=8/6). Direct comparison of infarct between males and females revealed no statistical difference with vehicle treatment, while demonstrating a sex-specific difference in the effect of capsinoid treatment (total infarct: two-tailed t-test, p=0.0447; percent infarct: two-tailed t-test, p=0.0216, n=8/8) (**Fig 2H-K**). These findings indicate a greater benefit of capsinoid-induced hypothermia in male mice.

**Figure 2:**
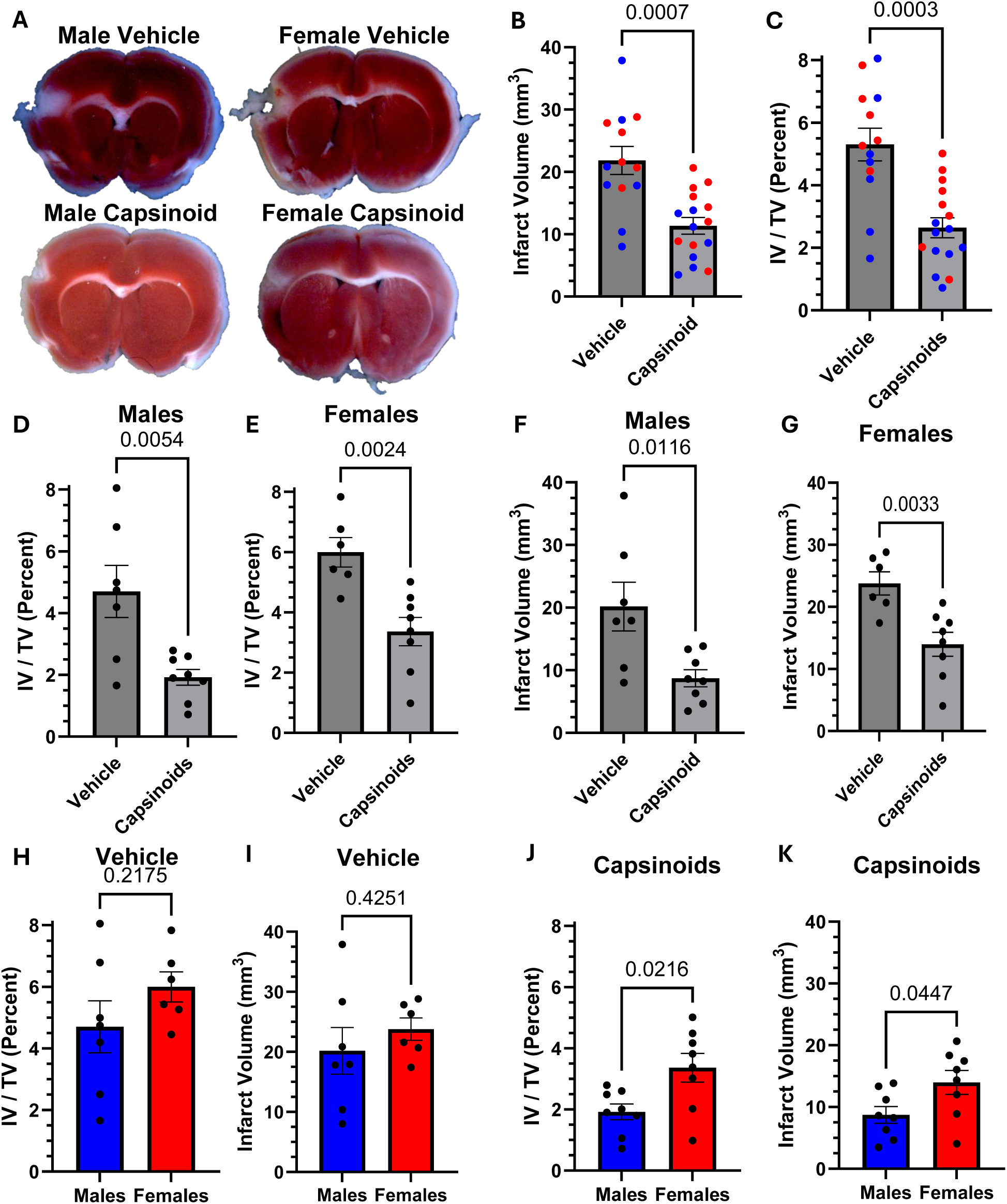
Capsinoid-induced hypothermia provides significant neuroprotection 3 days after pdMCAO. **A:** Representative images of TTC stained brains 3 days after pdMCAO separated by experimental condition and sex; white areas in the cortex represent infarct, red regions represent healthy tissue. **B:** Vehicle vs capsinoid comparisons of TTC Infarct volume (IV) divided by total volume (TV) represented as a percent (IV/TV) with males (blue points) and females (red points) combined. **C:** Vehicle vs capsinoid comparisons of TTC total infarct volume (in mm^3^) of males (blue points) and females (red points) combined. **D-G:** Vehicle vs capsinoid comparison of TTC IV/TV in males alone (**D**) and females alone (**E**) as well as total IV in males alone (**F**) and females alone (**G**). **H-K:** Male vs female comparisons of vehicle IV/TV (**H**) and total IV (**I**) and capsinoid IV/TV (**J**) and total IV (**K**). (p values presented as numbers)

To additionally evaluate the effect of capsinoid-induced hypothermia in a stroke model with reperfusion, we used the transient middle cerebral artery occlusion/reperfusion (MCAO/R) model. The MCAO/R transiently occludes the MCA at its origin, thus generating a larger infarct territory compared to the pdMCAO. The MCAO/R also models the reperfusion component that occurs with recanalization, such as with successful thrombectomy. Pilot studies revealed that the aged MCAO/R mice were not able to reliably maintain Tb at the end of the 2 hour heat support period. However, by 4 hours after stroke, MCAO/R mice were able to maintain Tb without ongoing heat support. Thus, for the MCAO/R studies, we used a 60 min occlusion with 4 hours of post-stroke heat support before administering either capsinoids or vehicle (**Fig 3A**). The baseline Tb measured during the last 30 minutes with heat support was similar between the treatment groups (Capsinoid: 35.2±0.2°C, Vehicle: 35.3±0.2°C, two-tailed t-test, p=0.6502, n=10/12). Capsinoid treatment induced a significant reduction in Tb (-4.3±0.4°C) compared to vehicle controls (-0.5±0.2°C) (**Fig 3B**, two-way RM measures ANOVA, p<0.0001, n=10/12). Notably, the capsinoid-treated mice demonstrated significantly better survival rate compared to vehicle-treated mice (**Fig 3C**, Capsinoid: 80.0% survival, Vehicle: 33.3% survival, Mantel-Cox, p=0.0272, n=10/12). Due to the high mortality in the vehicle treated mice after MCAO/R compared with the high survival in the pdMCAO model, we proceeded with the pdMCAO model for subsequent experiments examining the chronic stroke phase.

**Figure 3:**
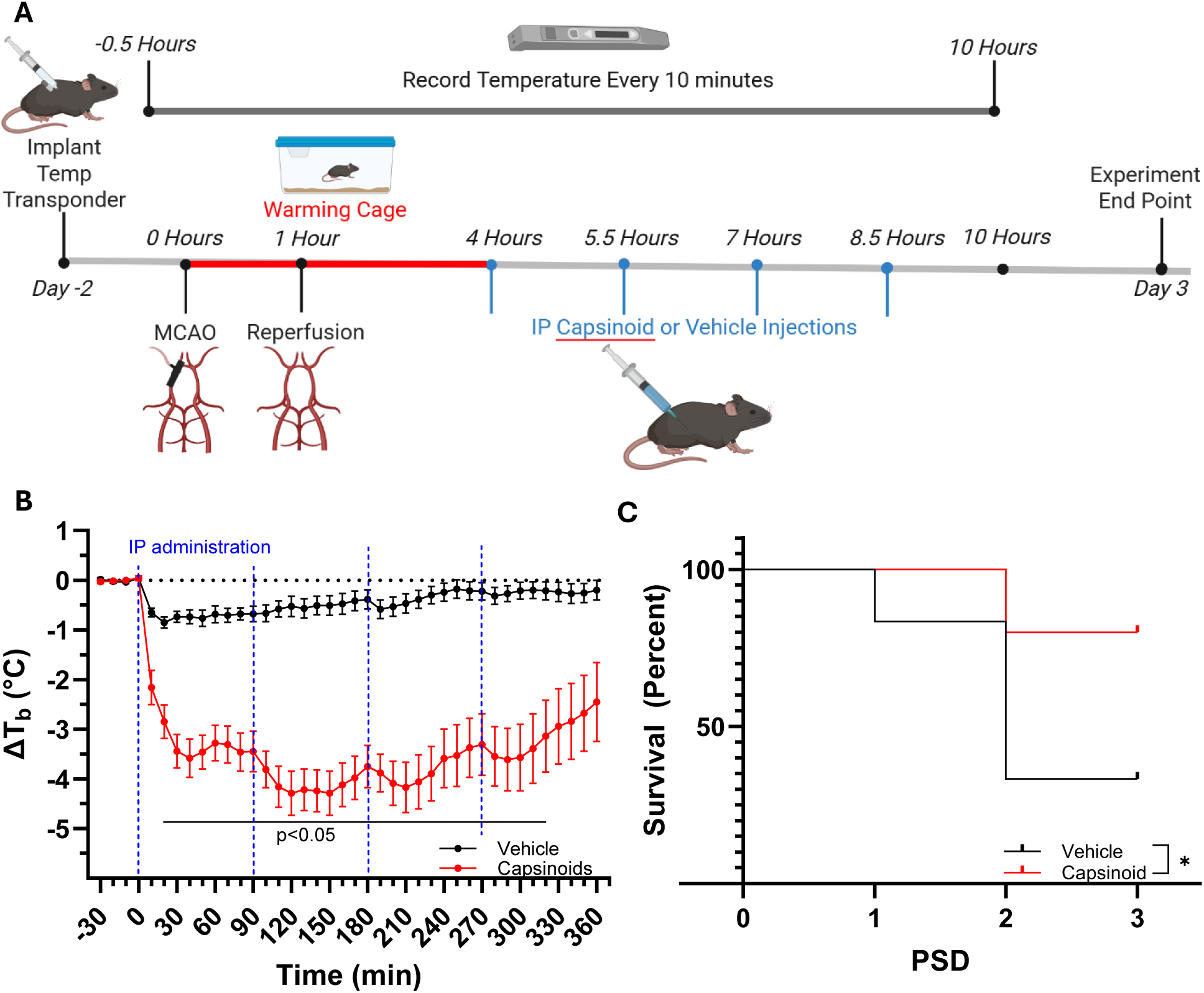
Capsinoid-induced hypothermia improves survival after MCAO/R. **A:** Experimental design showing timing of transponder implantation, MCAO/R occlusion and reperfusion points, recovery time, IP administration of vehicle or capsinoids, recording time, and experimental endpoints. **B:** Measured ΔTb of aged mice after MCAO/R stroke and repeated IP administration of capsinoids (40 mg/kg; indicated by blue dashed vertical lines) presented as the combined group of males and females. **C:** Survival plot comparing vehicle and capsinoid treated mice after MCAO/R up to 3 post-stroke days (PSD) with males and females combined.

Having established the benefit of capsinoid-induced hypothermia on acute stroke outcome, we next sought to determine if capsinoid treatment could similarly provide long-term improvement. For this purpose, we used the pdMCAO model for its consistent infarct size and high survival rate with aged mice. Capsinoid-mediated hypothermia (or vehicle/normothermia) was initiated at two hours after distal MCA occlusion (**Fig 4A**).

**Figure 4:**
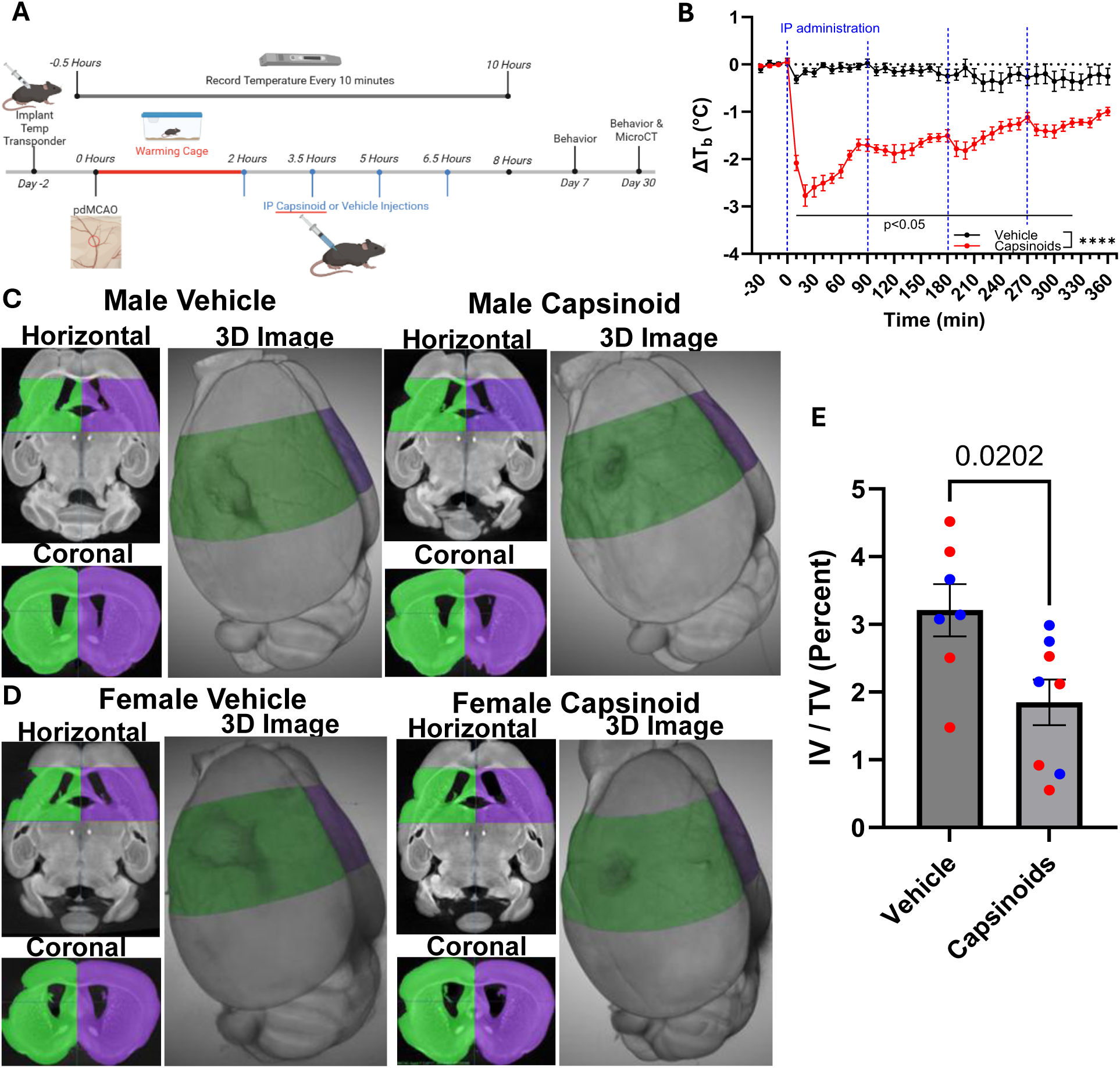
Capsinoid-induced hypothermia provides long-term histological neuroprotection. **A:** Experimental design showing timing of transponder implantation, pdMCAO stroke, recovery time, IP administration of vehicle or capsinoids, recording time, behavior tests at 1 week and 1 month, and experimental endpoints. **B:** Measured ΔTb of aged mice after pdMCAO stroke and repeated IP administration of capsinoids (40 mg/kg; indicated by blue dashed vertical lines) presented as the combined group of males and females. **C-D:** Representative images of microCT scans comparing vehicle and capsinoid treatments in horizontal and coronal views as well as 3D reconstruction of males (**C**) and females (**D**); ipsilateral hemisphere is colored in green and contralateral is colored in purple. **E:** Vehicle vs capsinoid comparison of microCT IV/TV with males (blue points) and females (red points) combined. (*: p<0.05, **: p<0.01, ***: p<0.001, #: p<0.0001, or listed as numbers)

Behavior tests were then performed to evaluate sensorimotor deficits at one week and one month post-stroke. Baseline Tb measured during the last 30 minutes in the warming cage (90-120 minutes after stroke onset) were similar between treatment groups (capsinoid: 36.9±0.1°C, vehicle: 36.8±0.3°C, two-tailed t-test, p=0.7456, n=8/7).

As with the acute cohort, the capsinoid-treated group demonstrated a significant decrease in Tb compared to vehicle controls (-2.8±0.2°C vs. -0.1±0.1°C, two-tailed t-test, p<0.0001, n=8/7) (**Fig 4B**). At one month post-stroke, brains were treated with iodine and examined by microCT (uCT) to quantify brain injury as tissue loss (**Fig 4C/D**). Ipsilateral and contralateral hemisphere volumes from a region encompassing the stroke injury were compared. Total tissue loss was determined by the difference between contralateral and ipsilateral volumes and expressed as a percentage of the total volume. Capsinoid treated mice demonstrated significantly less brain tissue loss (1.8±0.3%) compared with vehicle control (3.2±0.4%) (**Fig 4E**, two-tailed t-test, p=0.0202, n=8/7).

As the pdMCAO model predominantly induces injury within the sensorimotor cortex, we selected behavioral tests that evaluate function regulated by the same region. At one week and one month post-stroke, we tested mice for gait patterns, grip strength, and foot fault. Evaluation of gait characteristics at one week post stroke revealed capsinoid treated mice had a significantly shorter stance time (0.22±0.01s) in their left forepaw compared to vehicle treated mice (0.26±0.02s), indicating reduced weakness (i.e. better ability to keep up with the treadmill) in the contralateral limb of capsinoid-treated mice (**Fig 5A**, two-tailed t-test, p=0.0349, n=7/8). Additionally, capsinoid treated mice showed reduced variability in hind limb stride width compared with vehicle treated (**Fig 5B**) (0.22±0.02cm vs. 0.29±0.02cm, two-tailed t-test, p=0.0209, n=7/8) and significantly less gait symmetry loss compared to vehicle treated mice (**Fig 5C**) (1.00±0.01 vs. 0.95±0.02, two-tailed t-test, p=0.0302, n=7/8). In the latter case, the loss of gait symmetry (fore limb to hind limb) reflected an increase in hind limb stepping frequency to make up for a weaker step. Other measures of gait characteristics (swing, stride, propel times, stride frequency, stride length, or stance width) showed no significant differences between treatment groups.

**Figure 5:**
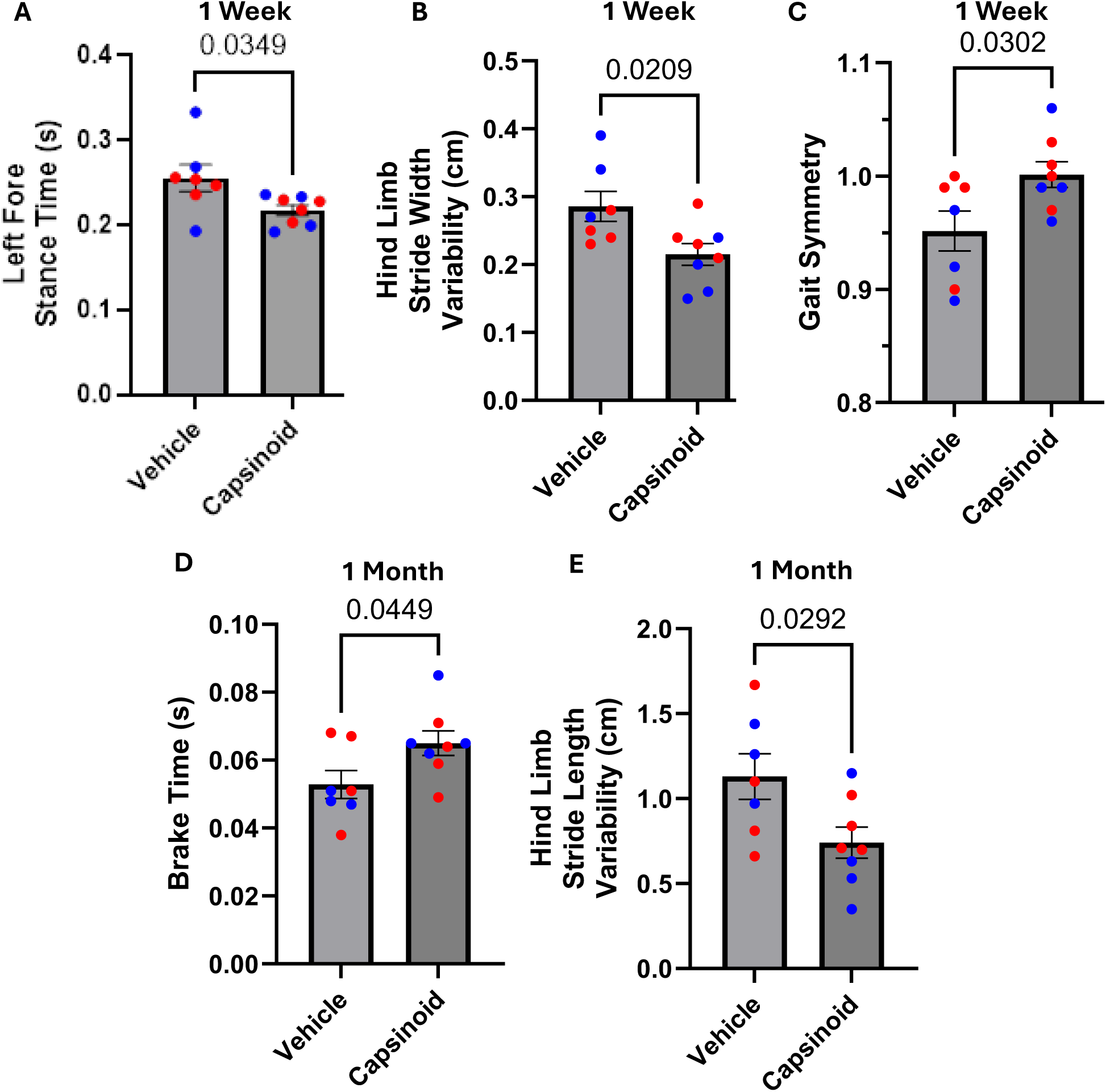
Capsinoid-induced hypothermia ameliorates long-term stroke induced motor deficits that present as gait abnormalities. **A-E:** DigiGait indices comparing vehicle and capsinoid treatment: 1 week left fore stance time (**A**), 1 week hind limb stance width variability (**B**), 1 week gait symmetry (**C**), 1 month left hind brake time (**D**), 1 month hind limb stance length variability (**E**). (p values presented as numbers)

By one month post-stroke, the gait differences described above were no longer significant, however, significant differences became evident in other gait measures. At this later time point, capsinoid treated mice (0.065±0.04s) had shorter left hind paw brake times compared with vehicle treated mice (0.053±0.004s), reflective of better surety in paw placement (**Fig 5D**, two-tailed t-test, p=0.0449, n=7/8). Additionally, hind limb stride length variability was significantly lower in capsinoid-treated mice (0.74±0.09cm) compared with vehicle treated mice (1.13±0.13cm) (**Fig 5E**, two-tailed t-test, p=0.0292, n=7/8). This reduced variability in hind limb stride parameters (stride width at 1 week and stride length at 1 month) are indicative of less motor weakness in capsinoid treated mice. In combination, significant differences in these gait indices indicate that capsinoid treated mice have reduced motor weakness in the contralateral limbs.

We also investigated the effects of capsinoid-induced therapeutic hypothermia by other measures of motor deficit, including grip strength and foot fault tests. Capsinoid treated mice showed significantly greater grip strength compared to vehicle treated mice at both one week and one month post-stroke (**Fig 6A**). Normalized grip strength at one week post-stroke was 0.91±0.04 versus 0.73±0.06 for capsinoid and vehicle, respectively (two-way RM measures ANOVA, p=0.0187, n=7/8). At one month, normalized grip strength was 0.96±0.05 versus 0.65±0.05 for capsinoid and vehicle, respectively (two-way RM measures ANOVA, p=0.0004, n=7/8). The foot fault test also revealed improved outcome with capsinoid-induced hypothermia. Contralateral foot fault percent was significantly less in capsinoid treated mice at both one week and one month following stroke (**Fig 6C**). Contralateral foot fault percent for capsinoid versus vehicle was 6.6±0.5 vs. 8.7±0.3% at one week (two-way RM ANOVA, p=0.0049, n=7/8) and 7.7±0.5 vs. 9.5±0.7% at one month (two-way RM ANOVA, p=0.0006, n=7/8). The total number of steps during the two minute trial did not differ by capsinoid or vehicle treatment at either one week (238.5±16.2 vs. 233.7±16.5; two-way RM ANOVA, p=0.8726) or one month (237.3±16.7 vs. 234.0±13.0; two-way RM ANOVA, p=0.8801) (**Fig 6B**). Further, when comparing the relative frequency of the contralateral foot faults to total foot faults, capsinoid treated mice demonstrated reduced frequency of contralateral foot fault at both one week and one month post stroke (two-way RM ANOVA, p<0.0001 and p<0.0001, n=7/8) (**Fig 6D**).

**Figure 6:**
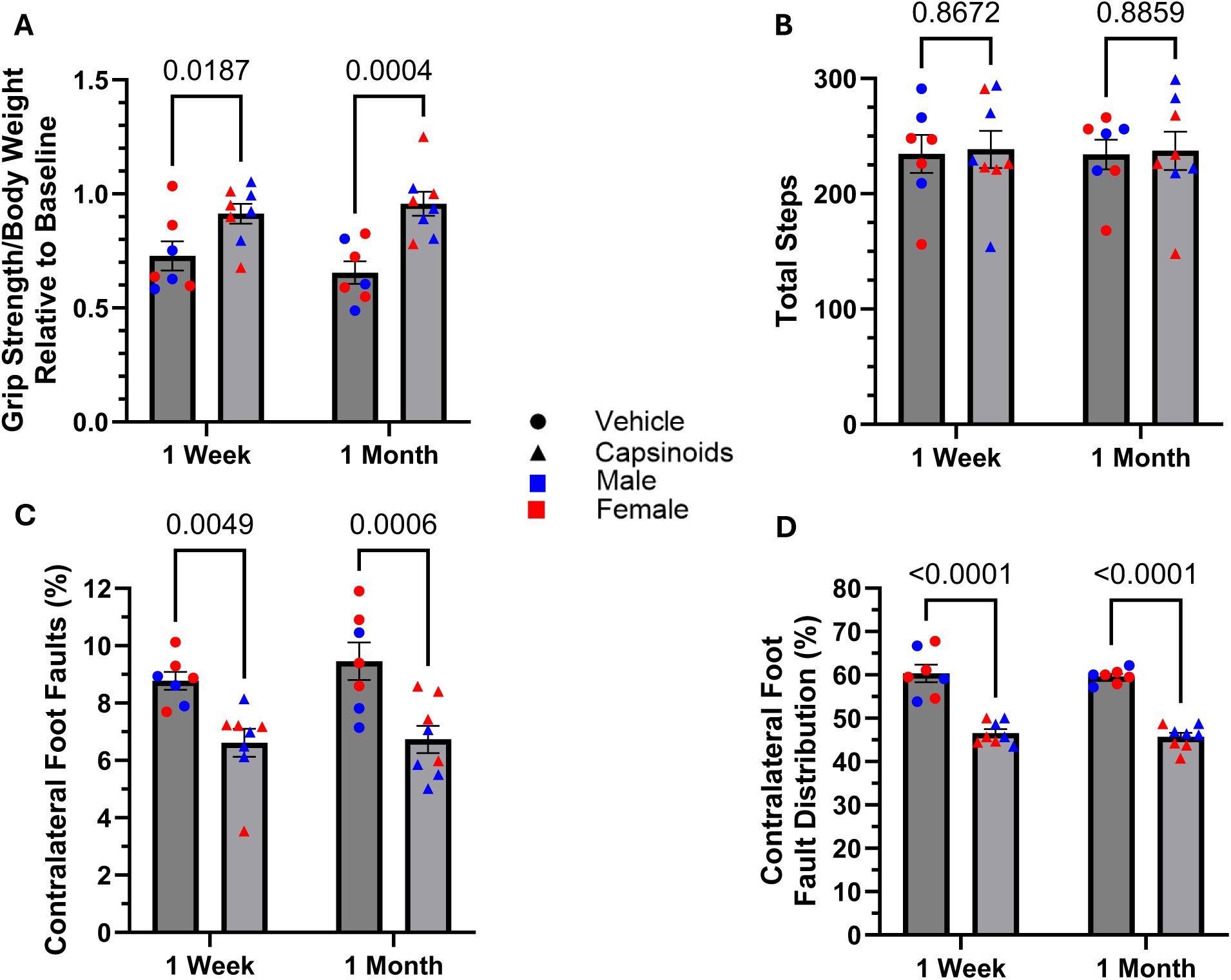
Capsinoid-induced hypothermia ameliorates long-term stroke induced sensorimotor deficit. **A-D:** Vehicle (circles) vs capsinoids (triangles) comparisons of grip strength normalized to both body weight and baseline (pre-stroke) values (**A**), contralateral foot faults as a percentage of total steps (**B**), total steps during foot fault test (**C**), and contralateral foot faults as a percentage of total foot faults (**D**) at 1 week and 1 month post-stroke with males (blue points) and females (red points) combined. (p values presented as numbers)

## Discussion

Therapeutic hypothermia (TH) has become the standard of care for ischemic brain injuries resulting from cardiac arrest and nHIE [15–24, 36, 52–61]. TH has also demonstrated considerable promise in experimental models of stroke. However, despite these successes in the laboratory, clinical trials have yet to produce convincing benefit for adult stroke patients [34, 36–38, 62]. Some limitations for successful application of TH in stroke patients relate to practical challenges in promoting body cooling in the awake patient. Current best methods of inducing hypothermia require heavy sedatives or paralytics to prevent violent shivering, and for those requiring paralytics, there is also the increased risk of ventilator-associated pneumonia [32, 35, 38, 63–69]. Most stroke patients are conscious throughout treatment and there are treatment advantages to maintaining their conscious state, such as for monitoring neurofunction. As such, improved methods of inducing hypothermia that are effective and compatible with the conscious patient are needed. Our previous studies have shown that pharmacological activation of TRPV1 channels by capsaicinoids (e.g. capsaicin and dihydrocapsaicin) can promote sustainable hypothermia and efficient suppression of the cold defense mechanism in conscious mice[31–33, 39–41]. Further, we showed that dihydrocapsaicin-mediated hypothermia can provide neuroprotection following stroke[31–33]. However, some studies indicate that TRPV1 channel activation can contribute to the ischemic injury mechanism and that activation of TRPV1 channels within the brain can exacerbate brain injury [70, 71]. Thus, an approach in which hypothermia is induced via the restricted activation of peripheral TRPV1 channels might spare the brain of potentially damaging neuronal toxicity. We recently showed that capsinoids (non-pungent TRPV1 agonists) can be effectively restricted to the peritoneal cavity due to the high lability of these molecules resulting from esterase vulnerability [46]. Further, peritoneal delivery of these capsinoids provided sustained hypothermia and neuroprotection in young mice. Here, we show that capsinoid mediated hypothermia provides improved outcome when initiated after stroke in two experimental stroke models using translationally relevant aged mice of both sexes. These studies used aged mice of both sexes to demonstrate that *1*) capsinoid treatment in the post-stroke period can decrease core body temperature into the mild hypothermic range, *2*) capsinoid-induced hypothermia reduces infarct size (pdMCAO model) or mortality (MCAO model) in the subacute phase after stroke, and *3*) capsinoid-induced hypothermia reduces brain atrophy and improves long-term functional outcomes in the chronic stroke phase.

### Capsinoids induce effective and consistent mild hypothermia when administered after stroke

Given that pharmacological activation of TRPV1 channels in the thermoregulatory system can induce mild hypothermia without triggering the cold defense mechanism, this approach offers a promising avenue to rapidly and efficiently achieve whole body cooling of conscious subjects. We demonstrate that IP administration of capsinoids 2 to 4 hours following stroke onset induces a mild hypothermia in aged mice and that it can be maintained for at least 6 hours. Capsinoid treatment induces a rapid temperature drop, triggering an initial drop in Tb within minutes and achieving target temperature within the mild hypothermia range within 10-20 minutes. We also show that neither the hypothermic response nor the beneficial effect of hypothermia was sex dependent.

Additionally, at the cessation of capsinoid treatment, the mice returned to normothermic levels spontaneously (normothermic in <2 hours after the final administration). The hypothermic response was not sex dependent.

### Capsinoid-induced hypothermia provides short-term neuroprotection after stroke in a translationally relevant aged cohort

Stroke is a disease that predominantly affects the elderly, with the majority of those affected being over 65 years of age[50, 72–75]. Compared to younger patients, older adults also face greater challenges in stroke recovery, including elevated mortality rates and more pronounced long-term impairments[50, 74, 76, 77]. Animal models of stroke also show mechanistic differences when comparing young to aged animals [78–81].

However, despite the clear role of aging in stroke, the majority of preclinical studies use young animals. Therefore, to improve the translational value of the present studies, we used aged mice to evaluate the effects of capsinoid treatment on experimental stroke outcomes. Using two experimental stroke models, our results demonstrated that the capsinoid-mediated mild hypothermia dramatically reduces stroke volume and improves survival during the sub-acute stroke phase. These findings of improved outcome align with other studies of experimental stroke that demonstrate neuroprotection using other methods to induce hypothermia [25, 68, 82].

Time to treatment is another important variable to consider for translational relevance to the treatment of stroke. Based on studies using more conventional forced cooling methods to induce hypothermia, optimal neuroprotection requires that cooling be initiated as early as possible [68]. For our studies, we incorporated a delay between stroke induction and the initiation of hypothermia to account for a better real-world applicability. Initiation of hypothermia was performed at either 2 or 4 hours post-stroke for the pdMCAO or MCAO/R models, respectively. With the successful demonstration of neuroprotection or improved survival in this study, future studies can examine how long hypothermia could be delayed before benefit is lost. Importantly, one must also factor the risks of hypothermia, such as the increased risk of infection with prolonged hypothermia. Therefore, it would also be important to determine the shortest duration of capsinoid-mediated hypothermia that still provides improved stroke outcome, while minimizing the risk of post-stroke infection or other risks associated with prolonged whole-body cooling.

Sex is another important biological factor when evaluating the effects of capsinoids on stroke outcome. Aged females have worse outcomes after stroke than their male counterparts, experiencing more severe strokes and worse recovery, both in the clinic and with experimental animal models [49, 51, 78, 83]. Capsinoid-induced hypothermia provided significant neuroprotection in the total cohort and when results were disaggregated by sex. However, while capsinoid-mediated hypothermia reduced infarct volume in both sexes, the benefit was significantly greater in males. There are several potential rationales for this difference in hypothermia benefit after stroke between sexes, including differing basal inflammation prior to injury, as well as differing contributions of multitudinous cell death mechanisms after injury. Several studies have demonstrated that microglia in aged male show a more pro-inflammatory profile compared to females [84, 85]. Given that one of the main mechanisms of hypothermia-driven hypothermia is suppression of inflammation, males starting with a more active inflammatory profile would likely benefit from capsinoid-induced hypothermia to a greater degree, as we observed [86]. Additionally, the mechanisms of cell death in response to ischemic injury have been shown to be sexually dimorphic with females showing more caspase-mediated cell death and males demonstrating more caspase-independent cell death, specifically through poly(ADP-ribose) polymerase-1 (PARP-1) and neuronal nitric oxide synthase (nNOS) [87]. Inhibition of this caspase-independent mechanism of apoptotic cell death led to neuroprotection in adult male mice but exacerbated stroke damage in adult female mice [88]. Additionally, hypothermia has been shown to reduce expression of both cleaved caspase-3 and PARP-1 in endothelial cells after ischemia-reperfusion injury [89, 90]. Therefore, the relative benefit of capsinoid-induced hypothermia on each of these cell-death pathways warrants further exploration.

### Capsinoid-induced hypothermia improves long-term stroke outcomes

To determine whether the neuroprotective effects of capsinoid-induced hypothermia persist beyond the acute phase, brain morphological and functional outcomes were assessed one month after distal middle cerebral artery occlusion (pdMCAO). Quantitative microCT imaging revealed markedly reduced cortical atrophy in capsinoid-treated mice compared with vehicle controls. Owing to its high volumetric precision and contrast sensitivity, particularly with iodine enhancement, microCT provides robust quantification of chronic infarct and tissue loss. The reduction in infarct volume indicates durable protection of cortical integrity and suggests that the early benefits of hypothermia also translate into long-term preservation of brain structure. Interestingly, at this chronic time point, the sex differences in hypothermia benefit were no longer evident. At this time, it is not clear if the converging outcomes at the chronic time point reflect sex differences in the delayed injury mechanisms or recovery/repair mechanisms.

Functional assessment demonstrated that these reductions in brain atrophy were associated with enhanced recovery of motor and sensorimotor function. In DigiGait analysis, capsinoid-treated mice exhibited superior locomotor performance relative to vehicle controls. At one week post-stroke, treated animals displayed features consistent with improved limb coordination and reduced motor asymmetry, including shorter contralateral forelimb stance time, reduced stride width variability, and preserved gait symmetry. By one month, early gross motor deficits had largely resolved in both groups; however, capsinoid-treated mice maintained greater stride stability and longer brake time in the contralateral hind limb, reflecting better control of paw placement and fine motor precision. These evolving patterns could suggest that capsinoid-induced hypothermia facilitates both early compensation and sustained refinement of motor recovery.

Consistent with the DigiGait findings, body weight-normalized forelimb grip strength was significantly preserved in the capsinoid group. While both groups exhibited early post-stroke weakness, the vehicle-treated mice showed continued decline over one month, whereas capsinoid-treated mice demonstrated partial recovery. In the foot fault test, capsinoid-treated mice also demonstrated improvement over vehicle-treated mice, committing fewer contralateral foot placement errors at both one week and one month. This attenuation of motor deficit further supports a lasting functional benefit of capsinoid-mediated hypothermia.

### Study Limitations and Future Directions

Due to the high mortality rate with the more severe MCAO/R model in aged mice, particularly in the vehicle/normothermia group, further histological and behavioral analyses were not able to be performed. However, future studies examining brain and functional outcomes in the sub-acute and chronic phases with this model could increase survival by shortening the occlusion time.

Chronic studies showed reduction of brain atrophy and improved behavioral testing measures in the sex-aggregated analyses. However, the experiments were underpowered to determine if sex-specific effects existed in the behavior and brain atrophy measures. Future prospective studies would be warranted to determine if there are sex-based differences in the chronic measures of brain injury or functional recovery.

### Summary and Conclusion

We have shown that intraperitoneal administration of capsinoids reliably induces a rapid, sustained mild hypothermia in conscious aged mice of both sexes. Capsinoids are vulnerable to break down by ubiquitous esterases, which likely account for findings in our lab and others supporting the idea that their activity is largely confined to the location of their delivery. In the present application, by delivering capsinoids to the peritoneal space, the brain is spared of additional TRPV1 channel activation that might aggravate ischemic injury and thereby counter the beneficial effects of brain cooling.

Regarding the effect of capsinoids on stroke injury and outcome with the pdMCAO model, we showed that capsinoid-induced hypothermia provides reduced brain infarct, less brain atrophy, and better functional outcome when initiated at 2 hours after stroke onset. This benefit was evident in both males and females, both within the sub-acute and chronic stroke phases. Interestingly, within the sub-acute phase, the benefit of capsinoid-mediated hypothermia was greater in males, suggesting that males may have a greater hypothermia-sensitive component versus females at this time point. However, by one month, sex-dependent differences in brain atrophy were no longer evident. With the more severe MCAO/R model, we showed significant improvement in survival with capsinoid-mediated hypothermia when initiated at 4 hours after stroke induction.

Overall, the significant decreases in brain infarct and atrophy along with improved short and long-term functional outcomes, provides a promising foundation for further studies in the application of capsinoid-mediated hypothermia for stroke treatment.

## Acknowledgements

This project has been supported by several sources of funding including: AA: AHA23PRE1023201, TL1T003169, UL1TR003167; SM: NIH R01AG081942, NIH R56NS120709, NIH R56AG084130.

Research reported in this publication was supported in part by the National Center for Advancing Translational Sciences of the National Institutes of Health under Award Numbers TL1TR003169 and UL1TR003167. This research was also supported in part by the U.S. Department of Agriculture (USDA)-Agricultural Research Service project 6060-41000-015-000D.MicroCT imaging was supported by NIH S10OD030336 and performed through the MicroCT Imaging Facility at the McGovern Medical School at UTHealth. The content is solely the responsibility of the authors and does not necessarily represent the official views of the National Institutes of Health.

## Author Contributions

AA and SM conceived of the project and designed the experimental paradigm. AA performed the bulk of the experimentation and analysis with significant help and support from SK, TW, AD, and GK. CC and RJ developed the capsinoid extraction process and provided the samples for experimentation. AA and SM wrote and edited the manuscript for publication and made the figures. All authors reviewed the manuscript.

## Competing Interests Statement

AA, SK, TW, AD, GK and SM declare no competing interests. CC and RJ are coinventors on a patent for a method to produce enriched capsinoid material.

## Data Availability Statement

All data generated or analyzed during the current study is available from the corresponding author on reasonable request.

## Notes

### Competing Interest Statement

The authors have declared no competing interest.

